# txtools: an R package facilitating analysis of RNA modifications, structures, and interactions

**DOI:** 10.1101/2023.08.24.554738

**Authors:** Miguel Angel Garcia-Campos, Schraga Schwartz

## Abstract

We present txtools, an R package that enables the processing, analysis, and visualization of RNA-seq data at the nucleotide-level resolution, seamlessly integrating alignments to the genome with transcriptomic representation. txtools’ main inputs are BAM files and a transcriptome annotation, and the main output is a table, capturing mismatches, deletions, and the number of reads beginning and ending at each nucleotide in the transcriptomic space. txtools further facilitates downstream visualization and analyses. We showcase, using examples from the epitranscriptomic field, how a few calls to txtools functions can yield insightful and ready-to-publish results. txtools is of broad utility also in the context of structural mapping and RNA:protein interaction mapping. By providing a simple and intuitive framework, we believe that txtools will be a useful and convenient tool and pave the path for future discovery. txtools is available for installation from its GitHub repository at https://github.com/AngelCampos/txtools.

## Introduction

The most widespread use of RNA sequencing data is quantifying gene expression, which merely requires counting the number of sequencing reads overlapping each gene. However, in addition to the identity of the gene to which they map, sequencing reads store a wealth of additional information, to which we will refer as ‘read-outs’, including (1) the relative position of read start, (2) the relative position of read ends, (3) the nucleotide identity frequency, and (4) the deletion counts in comparison to a reference sequence. These ‘read-outs’ have been leveraged to retrieve information pertaining to diverse layers of the RNA metabolism, including (but not limited to) RNA structure, RNA modifications, and RNA binding proteins. While information regarding these layers is typically lost when an RNA is reverse-transcribed into cDNA, pre-treatment of an RNA with diverse chemistries, enzymes, and/or affinity reagents can allow such information to be also maintained upon reverse transcription and to be captured in the form of one or more of the four above described ‘read-outs’. For instance, a wide range of RNA modifications - including m6A, m1A, pseudouridine, m5C, and ac4C - are all detected on the basis of RT-truncations (leading to an accumulation of read starts) (Linder et al. 2015; Safra et al. 2017; Hussain et al. 2013), RT misincorporations (leading to mismatches) (Schaefer et al. 2009; Squires et al. 2012; Edelheit et al. 2013; Li et al. 2017; Safra et al. 2017; Liu et al. 2022) or cleavage events (giving rise to accumulations of both read-starts and ends) that these modifications induce (Birkedal et al. 2015; Marchand et al. 2016; Garcia-Campos et al. 2019), typically after treatment with specific chemistries or enzymes. Similarly, approaches for mapping RNA structure, such as DMS-seq, CIRS-seq, SHAPE-seq, and LASER-seq, all rely on chemistries that preferentially interact and modify unstructured RNAs leading to RT-induced misincorporations (Aviran et al. 2011; Incarnato et al. 2014; Umeyama and Ito 2018; Zinshteyn et al. 2019). Finally, CLIP-based approaches for mapping out interaction sites between RBPs and mRNA, or derivatives used to map interactions between modification-specific antibodies and mRNA, also culminate in RT-misincorporations and truncations (Van Nostrand et al. 2016; Körtel et al. 2021).

Thus, while quantification of gene expression levels requires recording only *counts* of reads at the *gene* level, detection, and quantification of the above-discussed regulatory levels require quantifying the above-described ‘read-outs’, at the *single-nucleotide* level. Several tools have been developed to date to supply such single-nucleotide readouts. A widely used computational tool to obtain this kind of data is the samtools’ *mpileup* utility (Danecek et al. 2021). mpileup is a very fast tool and outputs a long table with three main values per genomic position: the number of reads covering the position, read bases, and sequencing quality. Another similar tool is JACUSA2 (Piechotta et al. 2022), which deals mainly with the detection of single-nucleotide variants and RT-arrests in NGS data. JACUSA2 supports two modes of sample setups: single-sample mode, which identifies variants against a reference sequence; and paired-samples mode, which identifies variants comparing samples from two conditions. Its output is a table that sets a score for each variant and the number of read bases at each genomic position. Yet another tool is RNAframework (Incarnato et al. 2018), a comprehensive tool for analyzing data derived from an array of RNA structure and RNA modification detection assays. While these tools, along with others (Picardi and Pesole 2013), offer many advantages, including speed and memory consumption, they suffer from two key - and partially related - limitations.

One limitation is that these tools were designed to operate in a single ‘space’, i.e. either genomic or transcriptomic. For RNA-centered quantifications, choosing only a single of these spaces is limiting. On the one hand, it makes intuitive sense to consider the entirety of the genomic space for mapping purposes, to avoid erroneous mapping of reads to wrong transcripts, which can occur if the transcripts from which they originated are not represented in the transcriptome annotation. On the other hand, the functional unit from which the read originated is the transcript, and hence accurate interpretation of the read mapping patterns (such as misincorporation, or premature termination) as well as downstream analyses must be on the basis of the transcriptomic, rather than genomic, space. A second limitation of current tools is in their handling of paired-end reads. Sequencing both ends of RNA fragments can be synergistically more informative than single-ended counterparts. Consider the case of using premature RT-truncations for the purpose of modification detection (e.g. pseudouridine, via CMC-based profiling (Carlile et al. 2014; Lovejoy, Riordan, and Brown 2014; Schwartz et al. 2014; Li et al. 2015). If only a single, short (e.g. 40 nt) read from one end is sequenced, each such read informs that no RT-truncations occurred at any of the 39 3’ terminal nucleotides of the read. However, if a mate read - again 40 nt long - is available, but with an insert size of 150 nt with respect to the first read, this mate read not only informs that no-RT mutations had occurred anywhere along the 40 nt long of the sequenced part of the read but also anywhere within the 70 (unsequenced) nucleotides separating the sequenced parts of the first and second read. Thus, despite the absence of any sequence coverage within the interval between the two reads, the availability of paired-end reads allows drawing inferences with respect to modification status. As an additional example, some protocols for mapping mRNA modifications rely on modification-dependent cleavage by RNAses sensitive to the modification (Garcia-Campos et al. 2019; Zhang et al. 2019). Again, paired-end reads provide critical information - in this case about the absence of cleavage - along the entirety of the intervening, non-sequenced elements between the two sequenced pairs. Yet, to our knowledge available tools do not integrate information originating from each of the two reads, and hence do not allow capturing the full extent of information present in paired-end reads.

Here we present txtools, an R package facilitating genomic data analysis, exploration, and visualization at the single-nucleotide resolution. txtools’ main output and data structure contains transcriptome-wide nucleotide-resolution coverage and ‘read-outs’, while seamlessly connecting genomic and transcriptomic coordinate systems. We provide several vignettes, showcasing the ability of txtools to perform start-to-end analysis and visualization of different types of datasets, via a few short and streamlined lines of code.

## Software

### Core functionality

txtools is an R package, readily available for installation through its GitHub repository AngelCampos/txtools. It strongly relies on Bioconductor infrastructure, especially on the GenomicRanges and GenomicAlignments packages (Lawrence et al. 2013) to load and process alignments data. Its main output and work-horse is of data.table class (Dowle and Srinivasan 2021), a high-performance version of R’s base data.frame class. For its plotting capabilities, txtools makes use of the ggplot2 package (Wickham 2016).

Its core inputs are BAM files harboring genomically aligned reads, a transcriptome annotation in BED12/6 format, and - optionally - a genomic reference sequence. The key output of the package is a table summarizing each of the four above-defined ‘read-outs’ at each position within the provided transcriptome space. Data processing is performed in two steps, 1) converting genomic space alignments into transcriptomic space alignments in a manner explicitly taking into account read mates (rather than single-end reads) when available, and 2) summarizing the four above defined ‘read-outs’ into a single-nucleotide resolution transcriptome-space based data table (Figure 1, top-panel).

**Figure 1.**
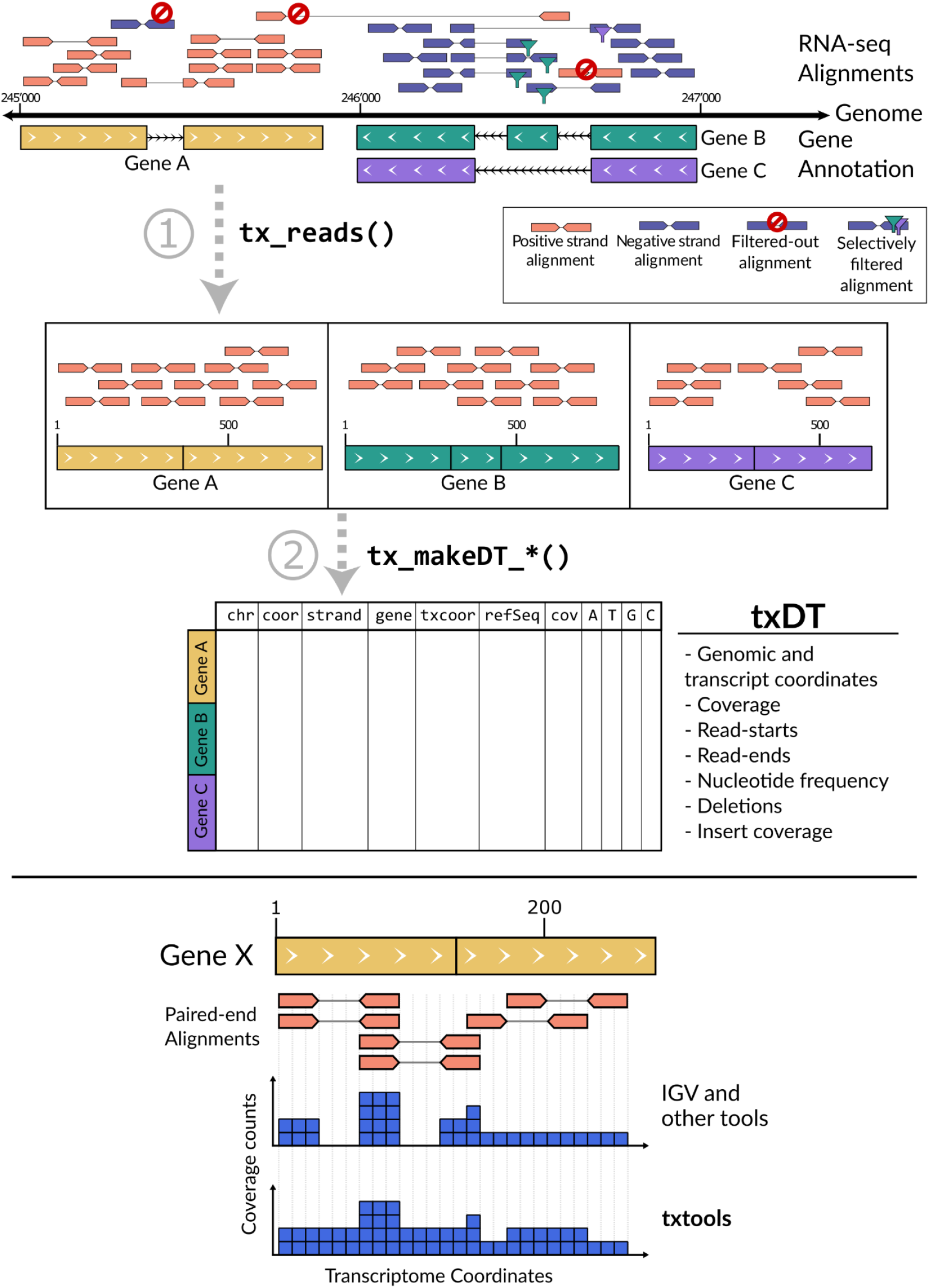
Top. txtools main RNA-seq reads processing. txtools allows easy loading of needed data: RNA-seq data in the form of reads mapped to a reference genome from BAM files, transcriptomic annotations in BED-12 or BED-6 format, and reference genomes from FASTA files. Using these data txtools can process the genomic reads into transcriptomic space, with tx_reads(), then generate a transcriptomic-wise summarized data table (txDT) using the tx_makeDT_*() family functions. txDT’s rows correspond to each nucleotide in a transcriptome, containing data on coverage, read-starts, read-ends, nucleotide frequency, and deletions. Bottom. Coverage counts in paired-end reads. Contrary to other tools that only count the sequenced stretch of a read towards coverage calculation, txtools counts the whole width of the paired-end read. This is from the start of read1 to the end of read2.

Processing of the genomic alignments into transcriptomic space is conducted with the tx_reads() function. This function implements filtering steps that will select only reads that are in the appropriate strand and within the gene structure, assigning all the reads that fit each gene’s (or isoform) boundaries defined in the gene annotation. Importantly, in the case of paired-end alignments read-1 and read-2 are merged into a single range, to allow coverage metrics to adequately consider the full insert, including the portion that is not sequenced between read-1 and read-2 (Figure 1, bottom-panel). txtools is able to process either single-end, paired-end reads, or long RNA reads, as well as those output by ONT sequencing technologies.

The second step consists of summarizing the transcriptomic reads into a table structure. This is performed through the tx_makeDT_* functions family, which has three options to summarize count data tables with 1) coverage, read-start, and read-end information; 2) only nucleotide frequencies and deletions; or 3) both data types. Both genomic and transcriptomic coordinate systems are provided in the summarizing table. If a genomic sequence is provided as input, the reference nucleotide at each transcriptomic position will also be reported. The data table provided as output uses the ‘data.table’ class from its eponymous package (Dowle et al. 2019), which offers enhanced functionality for large tables, compared to the base-R ‘data.frame’ class.

### Additional functionality

An array of additional functions are provided in txtools to facilitate common computational tasks required to analyze and visualize RNA-seq data at the single-nucleotide resolution. These include modules for calculating diverse metrics on the basis of the four read-outs, modules for sequence analyses, modules for metagene analyses, and ones for comparative, statistical analyses. Most of txtools’ functions have the prefix ‘tx_’ and are grouped by a following keyword:

- tx_load_*: Facilitate the input of files into the R session: The main input files for txtools consist of read alignments in BAM format (either paired-end or single-end), gene annotations in BED-12/6 formats, genome sequences in FASTA format, and previously saved txDTs in RDS format.
- tx_add_*: Add new information to an existing txDT, outputting the original txDT with the new column appended. Examples of these functions are tx_add_startRatio(), which adds the start to coverage ratio; tx_add_motifPresence(), which adds the location of RNA sequence motifs across the transcriptome; and tx_add_misincRateo(), which adds the ratio of misincorporation according to the genome and gene annotation. Importantly, these functions allow the seamless use of the pipe operator, from the *magrittr* package (Bache and Wickham 2014), enabling the chaining of one function after the other to create straightforward pipelines and easy-to-read code.
- *tx_get_*\*: Extract information from a txDT and generate an output that is not a txDT. These functions enable some downstream analysis and checks. Examples are *tx_get_flanksFromLogicAnnot*(), which extracts data from selected columns centering a window at positions specified by a logical variable, generating a matrix where each row represents a site of interest and each column a value in the window surrounding that site; another example is *tx_get_metageneRegions*() which outputs a metagene matrix with each row representing a gene and each column a bin in one of the codifying gene regions.
- *tx_plot_*\*: txtools plotting functions seek to fulfill two objectives: inspection of summarized data, and presentation of results. Examples are *tx_plot_nucFreq*() and *tx_plot_staEndCov*(), which plot the counts of data of nucleotide frequency, and read-starts/ends and coverage respectively; *tx_plot_seqlogo*(), which plots the sequence motif at sites annotated with a logical vector; and *tx_plot_metageneRegions*(), which creates metagene plots to compare values aggregated by gene regions.
- *tx_test_*\*: Use of the txDT objects from experimental data to compare metrics between groups. High-speed t-tests (Gentleman et al. 2019) and likelihood-ratio tests from edgeR (Robinson, McCarthy, and Smyth 2010) are implemented in txtools. These tests can be used to detect differences between groups in continuous variables in the first case or as ratios of count data in the second; for example, start to coverage ratio when testing for RT-stoppage in m6A detection using miCLIP.

## Use cases

The core txtools workflow for most projects would be as follows:

1. Load data into R: read alignments (BAM file), genomic annotation (BED12 format), and optionally a reference genome (FASTA).
2. Process alignments into txDTs.
3. Calculating metrics to support analyses: e.g. read-start to coverage rate, misincorporation rate, gene regions, etc.
4. Detect nucleotides that are relevant to the study.
5. Plot results.

Following we briefly present three use cases showcasing the ability of txtools to facilitate streamlined analysis of RNA-seq datasets at the nucleotide level. While the examples we chose are all in the context of detecting RNA modifications, similar workflows can be used for quantifying RNA structure and for analysis of CLIP-seq datasets.

### Case study 1: rRNA pseudouridylation detection

For the first use case, we analyzed pseudouridine mapping data, acquired via Pseudo-seq, a method for genome-wide, single-nucleotide resolution identification of pseudouridine (Carlile et al. 2014). Carlile and collaborators employed N-cyclohexyl-N′-(2-morpholinoethyl)-carbodiimide metho-p-toluenesulphonate (CMC) which reacts with pseudouridines creating an adduct that blocks cDNA’s reverse transcription one nucleotide downstream to the pseudouridylated site. The FASTQ data of a CMC-treated sample and a control sample were downloaded from GEO: GSE58200, and aligned to the yeast ribosome reference (Taoka et al. 2016) sequence using Rsubread (Liao, Smyth, and Shi 2019) to create the respective BAM files. Below are the steps and txtools functions used to analyze the data.

1. Load the reference genome with *tx_load_genome*(), and the gene annotation with *tx_load_bed*().
2. Process the resulting BAM files using the bam2txDT.R script, included in the txtools installation. This script processes the genomic alignments into transcriptomic ones using *tx_reads*() and generates a data table with summarized counts for coverage, read-starts, and read-ends using *tx_makeDT_coverage*().
3. Visualize the processed data with *tx_plot_staEndCov*() and observe the effect of CMC treatment on RT premature stoppage at a known pseudouridylated site, which manifests as an abrupt increase of read-starts compared to the control sample (**Figure 2A, top, and middle panel**).
4. Calculate the <u>s</u>tart-<u>r</u>ate <u>1bp d</u>own-<u>s</u>tream (SR_1bpDS), using *tx_add_startRatio1bpDS()* in both samples and subtract the resulting SR_1bpDS of the control from that of the CMC-treated sample, yielding the start-rate difference <u>1bp d</u>own-<u>s</u>tream (SRD_1bpDS), a metric that we can use to detect pseudouridines in CMC-treatment data.
5. Plot these results using the *tx_plot_numeric*() function to plot numeric variables in a txDT along a window in a gene (**Figure 2A, bottom panel**).
6. Using the resulting txDT and a common call to ggplot2’s scatterplot, the results for all four ribosomal transcripts are shown in **Figure 2B**. The resulting SRD_1bpDS metric shows that a simple threshold can discriminate between pseudouridine harboring and non-harboring sites on rRNA.

**Figure 2.**
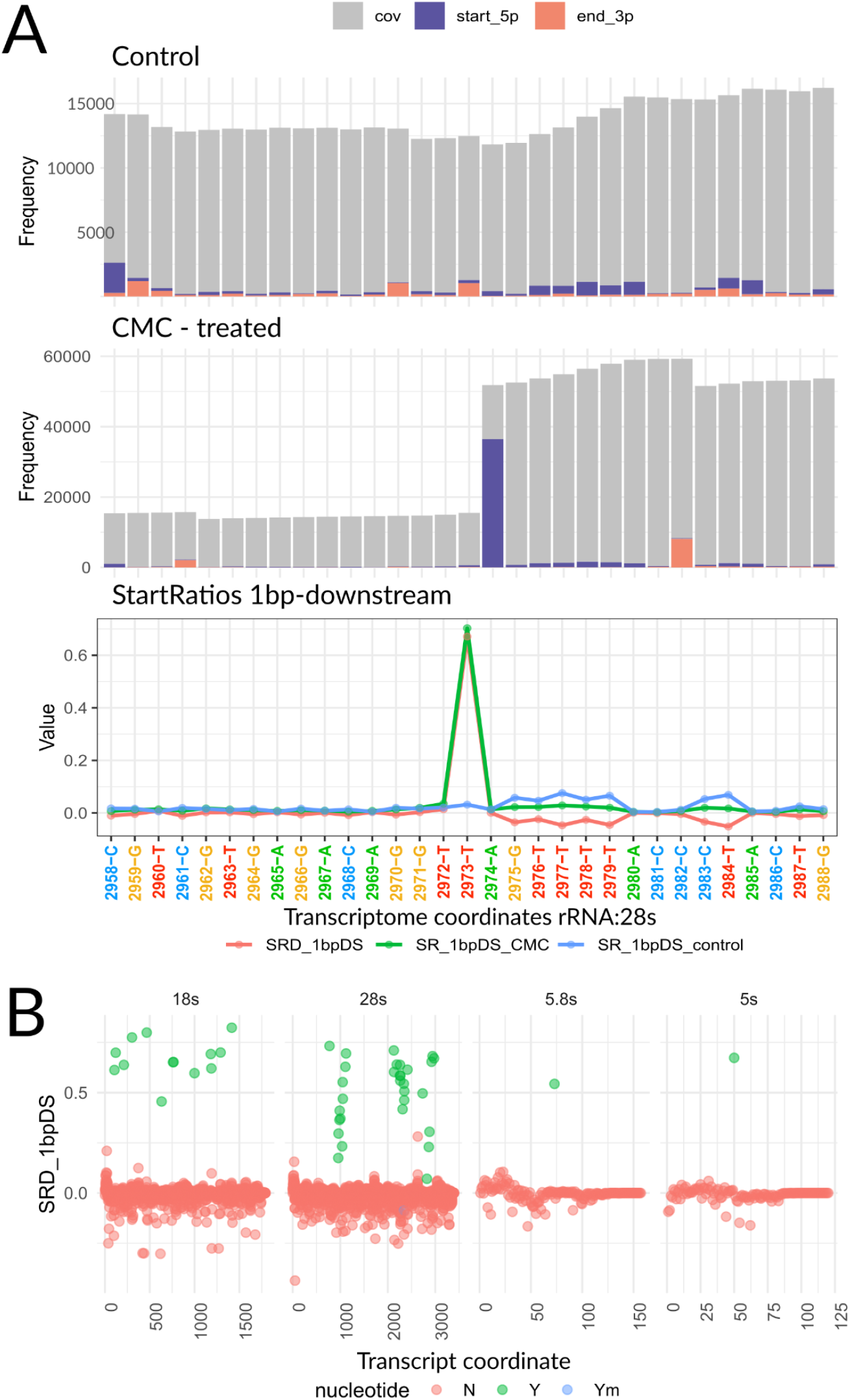
Use case #1 - rRNA pseudouridylation. A) Top. txtools Starts-Ends-Coverage plot for the mock-treatment sample centered at position 2973 of the yeast 28s rRNA, a known pseudouridine site. Middle. Same plot as above but for the CMC-treated sample. A dramatic difference in read-starts is evident at 1 bp downstream from the pseudouridylated site. Bottom. Lineplot showing SR_1bpDS for both CMC and control treatment and the resulting difference, SRD_1bpDS, at 28s:2973 and surrounding nucleotides. B) Scatterplots of full rRNA transcripts showing the SRD_1bpDS per nucleotide. Marked in green are the known pseudouridylated sites and marked in blue is the sole 2’-O-methylated pseudouridine.

### Case study 2: Single-nucleotide resolution detection of m6A using miCLIP2

For this case study we analyzed the data of the study that presented miCLIP2 (Körtel et al. 2021), a miCLIP enhancement allowing antibody-based single-nucleotide resolution mapping of m6A sites, relying on crosslinking of an antibody to methylated sites (Linder et al. 2015). Similarly to its predecessor, miCLIP2 relies on premature termination of reverse transcription during cDNA synthesis at the cross-linked residue. Data was downloaded from GEO: GSE163500, selecting for samples of wild-type (WT) and methyltransferase-like 3 (Mettl3) knockout in mESC, and subsequently aligned to the mm9 reference genome using Rsubread (Liao, Smyth, and Shi 2019). Below are the steps and txtools functions used to analyze the data.

1. Load the reference genome with *tx_load_genome*(), and the gene annotation with *tx_load_bed*().
2. Process all BAM files by looping through each with:
  a. *tx_load_bam*(): Loading the BAM file
  b. *tx_reads*(): Processing into transcriptomic reads
  c. *tx_makeDT_coverage*(): Processing into a table with summarized data on coverage, read-start, and read-ends.
  d. *tx_add_startRatio1bpDS*(): Adding the start ratio 1bp down-stream
3. Unify all the resulting tables with *tx_unifyTxDTL*(), this is to have them share the same transcriptomic coordinates, by selecting the intersection of genes in all data tables.
4. Perform t-tests using the txtools’ inbuilt ‘genefilter’ wrapper (Gentleman et al. 2013) *tx_test_ttests*(), testing for differences in startRatio_1bpDS between WT and KO samples both with the immunoprecipitation and crosslinking treatment.
5. Add DRACH motif presence with *tx_add_motifPresence*().
6. Plot a volcano plot using a call to ggplot2’s scatterplot, color-coding for DRACH presence, and select putative m6A sites at mean difference > 0.05 and p-value < 0.01 (**Figure 3A**).
7. Plot a metagene profile using *tx_plot_metageneRegions*(). Showing the previously reported enrichment of m6A sites just after the CDS end, compared to the baseline DRACH motif presence (**Figure 3B**).

**Figure 3.**
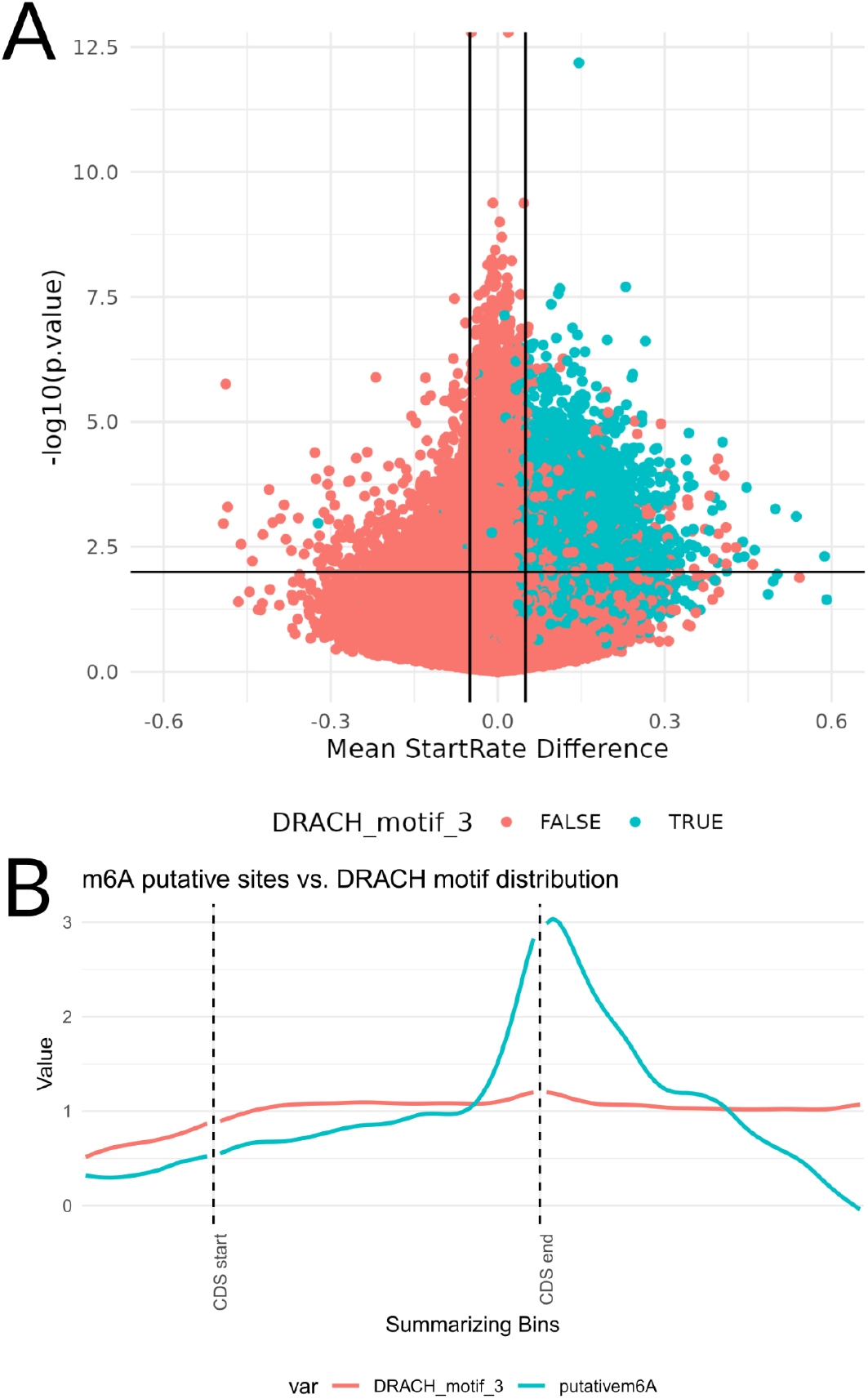
Case study #2 - m6A epitranscriptome in mESC using miCLIP2. A) Volcano plot showing the 1bp-down-stream mean start-rate difference at each queried position of the transcriptome (x-axis) and the -log10 p-value, calculated using a t-test (y-axis). Colored in blue are all sites that are centered in a DRACH motif. B) Metagene plot aligned at the end of CDS. Showing the relative abundance of putative m6A sites in blue, compared to the baseline presence of the DRACH motif in red. Putative sites thresholds used were startRatio_1bpDS difference > 0.05, p-value < 0.01, only considering adenines.

### Case study 3: Dynamic RNA acetylation using ac4C-seq

For this case study we analyzed RNA acetylation data, acquired via ac4C-seq, a chemical method for the transcriptome-wide quantitative mapping of N4-acetylcytidine (ac4C) at single-nucleotide resolution (Sas-Chen et al. 2020). Sas-Chen and collaborators employed the reaction of ac4C with sodium cyanoborohydride (NaCNBH3) under acidic conditions, which leads to C->T mutations at acetylated positions (Thomas et al. 2018). A key result in this study was the discovery of an abundance of acetylation sites on rRNA derived from the hyperthermophilic archaea *T. kodakarensis*. Data from GEO: GSE135826 was downloaded and aligned to the T. kodakarensis genome using Rsubread (Liao, Smyth, and Shi 2019). Below are the steps and txtools functions used to analyze the data.

1. Load the reference genome with *tx_load_genome*(), and the gene annotation with *tx_load_bed*().
2. Process the BAM files with the *bam2txDT*.*R* script provided along the txtools installation, with the parameters “-p TRUE -d covNuc -r 300 -m 0” to generate a summarized count data table with coverage, read-starts, read-ends, nucleotide frequency, and deletion frequency information.
3. Plot a nucleotide frequency plot using the *tx_plot_nucFreq*() function. To observe the high levels of misincorporations of cytidines in place of thymines exclusively in the NaCNBH3-treated samples (**Figure 4A**).
4. Calculate the C to T misincorporation rate across every position of the reference transcriptome with *tx_add_CtoTMR*(). and subtract the C to T misincorporation between the control and the NaCNBH3 treatment samples for each growth temperature to obtain the rate C to T misincorporation rate difference (MRD_CtoT).
5. Call putative ac4C sites if a position in any of the samples surpassed a threshold of MRD_CtoT > 0.005.
6. Plot a boxplot showing the calculated MRD_CtoT across temperatures (**Figure 4B**). The boxplot shows how the stoichiometry of ac4C sites increases as the temperature of growth increases, reaching the highest levels at 95 °C.

**Figure 4.**
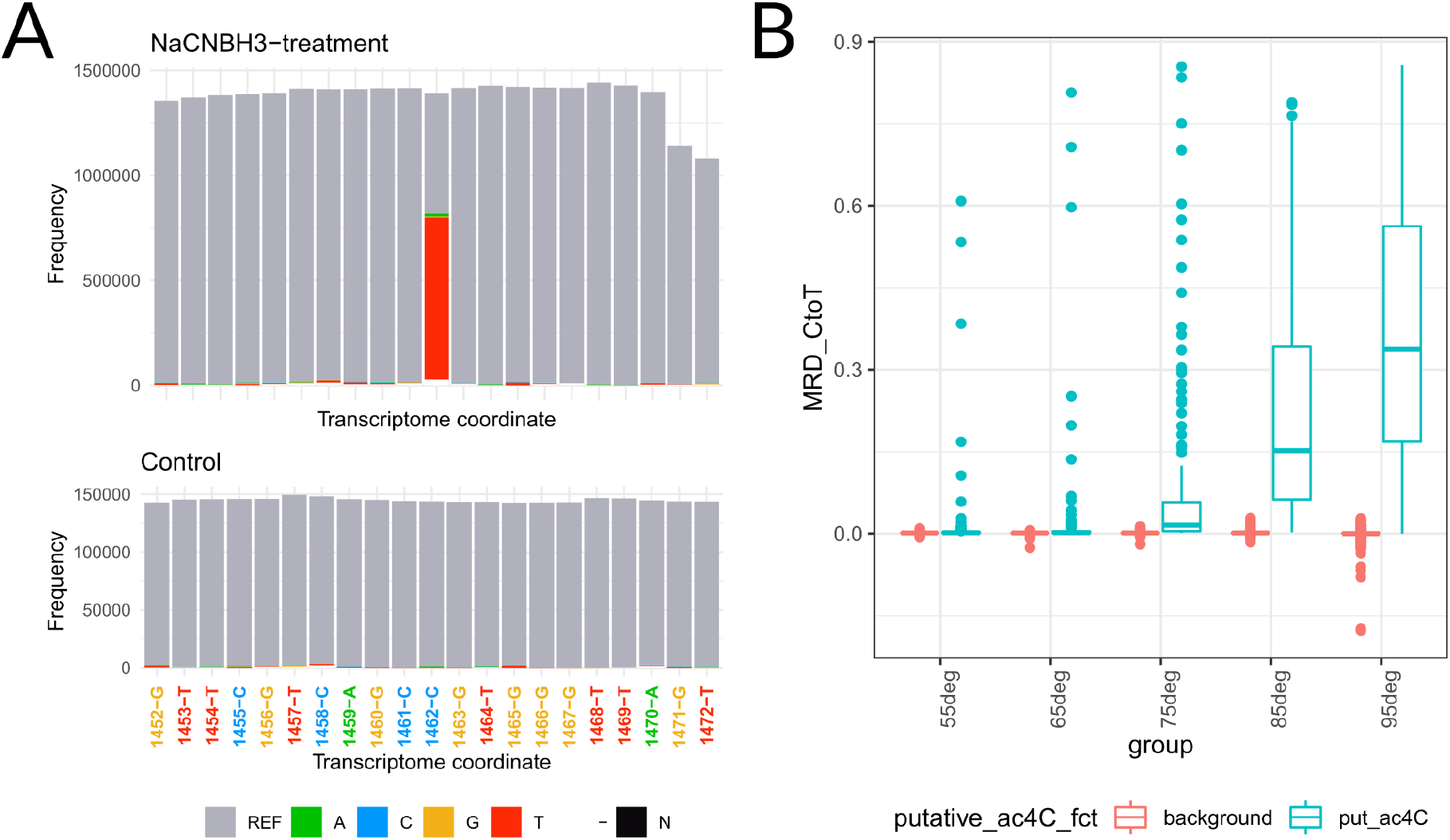
Case study # 3 - Dynamic RNA acetylation using ac4C-seq. T. kodakarensis’ ac4C levels increase across a temperature gradient. A) txtools’ nucleotide frequency plot. B) Boxplot of MRD_CtoT at putative ac4C sites at increasing growth temperatures.

Code to reproduce the three study cases presented is available at: https://github.com/AngelCampos/txtools_uc.

## Limitations

A current limitation of txtools is the potentially long processing times. To ameliorate this issue, txtools provides the ability to run many of its functions using multiple cores/threads taking advantage of the R package *parallel* (Team 2023) reducing the processing time. Of note, this capability is available only in UNIX-based systems and is not available in Windows systems. To provide a guide and expectation of the time taken to process data in different scenarios we arranged a set of benchmarks, using the hg19 genome and gene annotation to generate artificial paired-end reads. All scenarios simulate paired-end RNA-seq reads of 50 bp long for each read1 and read2 and an insert of 100 bp between the reads, extracted from the hg19 transcriptome which were then aligned to the genome using STAR (Dobin et al. 2013). The resulting BAM files were subject to four tasks:

- T1 = Converting RNA-seq genomic ranges into transcriptomic ranges without their nucleotide sequence.
- T2 = Generating a summarized table with coverage, read-starts, and read-ends data.
- T3 = Converting RNA-seq genomic ranges into transcriptomic ranges including their nucleotide sequence.
- T4 = Generating a summarized table with nucleotide frequency, insert coverage, and deletion frequency, as well as coverage, read-starts, and read-ends data.

Both T1 and T3 are performed by the function *tx_reads*(), T2 by *tx_makeDT_coverage*(), and T4 by *tx_makeDT_covNucFreq*(). In a common txtools processing workflow T1 will be followed by T2, and T3 by T4 to obtain a summarized count-data table.

The four tasks were carried out under different scenarios: 1) doubling the number of genes in the gene annotation from 2,500 up to 20,000 genes (Figure 5A), 2) doubling the number of processed paired-end reads from 4 million to 32 million (Figure 5B), 3) increasing the number of cores from 4, 8, to 12 (Figure 5C), and 4) increasing both the number of cores and the number of genes (Figure 5D), to show the time taken by the main processing functions of txtools. All tasks were run 5 times for each benchmark.

**Figure 5.**
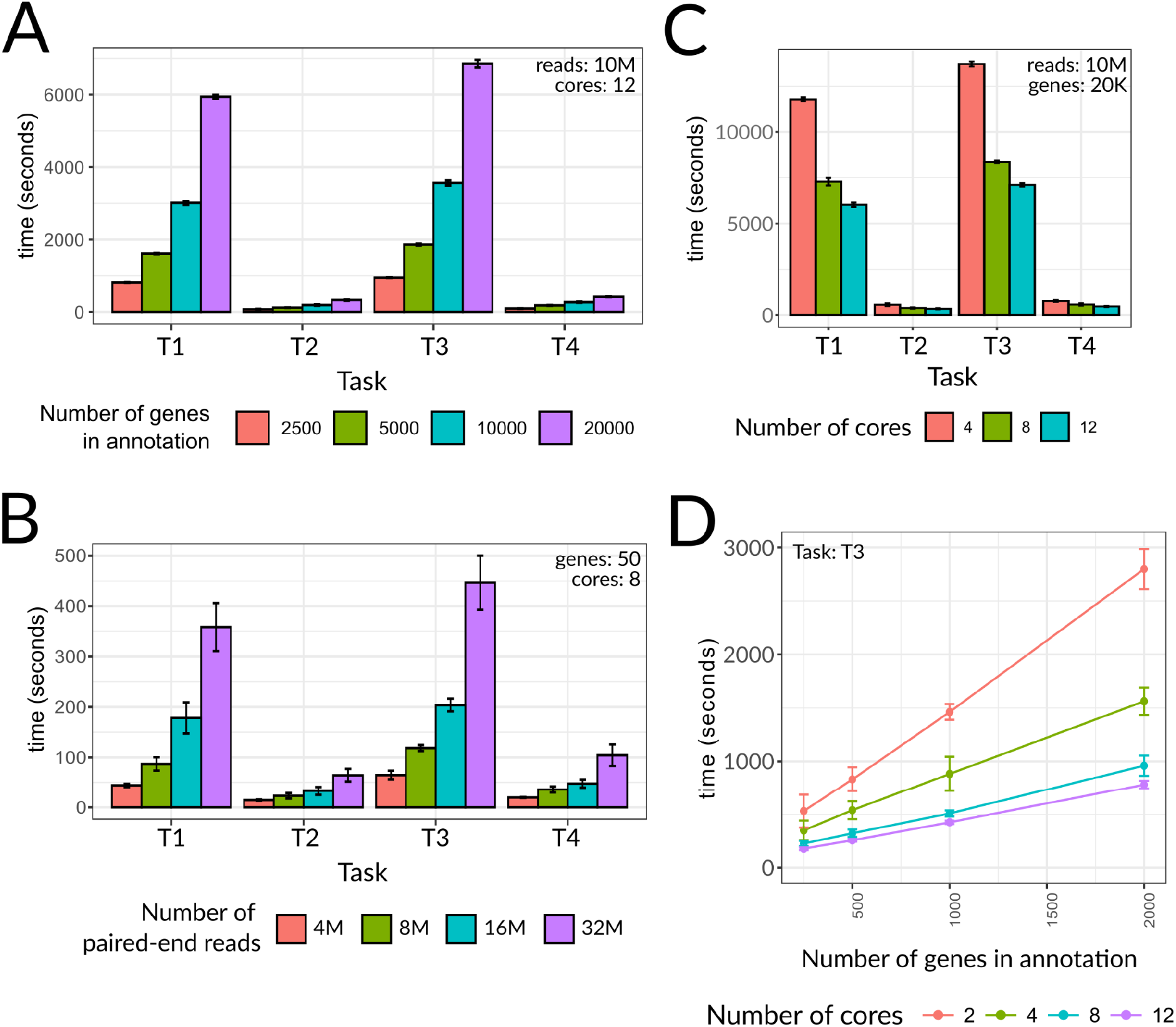
Processing time of the main txtools processing functions for different simulated scenarios. A) Barplot showing the processing time (y-axis) to perform tasks 1 through 4 (x-axis) in doubling numbers of genes (color-coded). B) Barplot showing the processing time (y-axis) to perform the benchmarking tasks (x-axis) in doubling numbers of simulated RNA-seq paired-end reads (color-coded). C) Barplot showing the processing time (y-axis) to perform the benchmarking tasks (x-axis) using 4, 8, and 12 cores (color-coded). D) Lineplot showing the processing time (y-axis) to perform task “T3” using 2, 4, 8, and 12 cores and doubling the number of genes (x-axis). Additional parameters used for each scenario are written in the top corner of each plot. M= Millions; K = Thousands. Each value represents the mean of 5 repeats and whiskers show the standard deviation.

The benchmarks show that the lengthier tasks are T1 and T3, processing the genomic reads into transcriptomic merging each read1 with their respective read2, carried out by the *tx_reads*() function. T3 is only slightly lengthier than T1, by adding the nucleotide sequence which carries a slight overhead. On the other hand, T2 and T4, which summarize the transcriptomic reads into count-data tables, carried out by *tx_makeDT*\*() functions, are much less time-consuming in comparison, and again with a slight overhead by incorporating nucleotide sequence information.

In terms of processing times, these scale roughly linearly with both the number of genes in the gene annotation and the number of reads/alignments in the simulated RNA-seq library (Figure 5A-B). On the other hand, running any task using an increasing number of cores also reduces the processing time in all tasks (Figures 5C-D). Nevertheless, the extent of this effect is not linear as spreading the process across multiple cores has the overhead of collecting and integrating the resulting outputs. In the scenario of Figure 5D doubling the number of cores from two to four reduced the processing time by 45% while doubling from 4 to 8 further reduced it by an additional ∼40%.

## Code availability

txtools source code is available from its GitHub repository at: https://github.com/AngelCampos/txtools. A manual for all txtools functions and vignettes showing examples of their functionality is available at the txtools package website. Links to the use cases and benchmarking code are also available through the txtools GitHub page.

## Discussion

txtools facilitates the processing and analysis of RNA-seq datasets at the single-nucleotide resolution level. The illustrated case studies showcase the power of txtools, and its ability to streamline analyses that are otherwise non-trivial to implement. txtools offers unique advantages in the processing of paired-end reads, where it allows quantifying information that is not quantified by any other tool, to our knowledge. The seamless integration between genomic and transcriptomic space is another unique advantage of txtools over additionally available tools. Finally, its availability as an R package allows it to readily be integrated into analytic workflows.

Additionally, to the case studies shown here, which rely on Illumina-based RNA-seq data, we have also successfully employed txtools for analysis of RNA-seq data from other platforms, such as Nanopore minION’s long reads.

A key limitation of txtools is its relatively long processing time. Processing time scales primarily with the number of genes and with the number of reads/alignments to be processed. Depending on needs and research focus, trimming down the annotated gene set can help substantially reduce running times. Another option is the txDT subsampling function *tx_sampleByGenes*(), which enables working only on a random sub-sample of the transcriptome. Once the analytic pipeline is more mature it can then be easily run with the full dataset to generate complete results. Additionally, to avoid long waiting times during an interactive R session, we offer the command line bam2txDT.R script that implements the core functionality of txtools and that can be run in the background. Nevertheless, we hope to optimize processing times in future versions of txtools.

We believe that txtools will be a useful and convenient tool for the RNA and bioinformatics communities. By providing a simple, intuitive, streamlined, and rich framework for processing, analyzing, and visualizing RNA-seq data at the nucleotide level, it paves the path for future discovery for both entry-level and advanced users.

